# Three-Dimensional Bioconjugated Liquid-Like Solid (LLS) Enhance Characterization of Solid Tumor - Chimeric Antigen Receptor T cell interactions

**DOI:** 10.1101/2023.02.17.529033

**Authors:** Duy T. Nguyen, Ruixuan Liu, Elizabeth Ogando-Rivas, Alfonso Pepe, Diego Pedro, Sadeem Qdasait, Nhi Tran Yen Nguyen, Julia M. Lavrador, Griffin R. Golde, Ryan A. Smolchek, John Ligon, Linchun Jin, Haipeng Tao, Alex Webber, Simon Phillpot, Duane A. Mitchell, Elias J Sayour, Jianping Huang, Paul Castillo, W. Gregory Sawyer

## Abstract

Cancer immunotherapy offers lifesaving treatments for cancers, but the lack of reliable preclinical models that could enable the mechanistic studies of tumor-immune interactions hampers the identification of new therapeutic strategies. We hypothesized 3D confined microchannels, formed by interstitial space between bio-conjugated liquid-like solids (LLS), enable CAR T dynamic locomotion within an immunosuppressive TME to carry out anti-tumor function. Murine CD70-specific CAR T cells cocultured with the CD70-expressing glioblastoma and osteosarcoma demonstrated efficient trafficking, infiltration, and killing of cancer cells. The anti-tumor activity was clearly captured via longterm in situ imaging and supported by upregulation of cytokines and chemokines including IFNg, CXCL9, CXCL10, CCL2, CCL3, and CCL4. Interestingly, target cancer cells, upon an immune attack, initiated an “immune escape” response by frantically invading the surrounding microenvironment. This phenomenon however was not observed for the wild-type tumor samples which remained intact and produced no relevant cytokine response. Single cells collection and transcriptomic profiling of CAR T cells at regions of interest revealed feasibility of identifying differential gene expression amongst the immune subpopulations. Complimentary 3D in vitro platforms are necessary to uncover cancer immune biology mechanisms, as emphasized by the significant roles of the TME and its heterogeneity.

## INTRODUCTION

Despite a better understanding of cancer biology and the discovery of new targeted cellular therapies, many patients still succumb to refractory solid cancers [1–4]. Chimeric antigen receptor (CAR) T therapy has shown great promise against hematological malignancies with complete remission rates as high as 90% during the first-month post CAR T cell infusion, leading to US FDA approval of now six CAR T-cells for B-cell malignancies [5–10]. These results however have not been translated to patients with solid tumors. The heavily immunosuppressive tumor microenvironment (TME) has presented barriers to CAR T cell trafficking, proliferation, and persistence in solid tumors [11–14]. Nevertheless, the mechanisms of immunosuppression in the solid TME remain poorly understood.

Approaches to address this knowledge gap include methods for CAR T cell delivery [15], engineering CAR T cells to improve target specificity and resist immunosuppressive factors in the TME [16], and reprograming the TME [17]. Before we can design more potent CAR T cells able to overcome the solid tumor TME, it is imperative to develop preclinical models to reliably evaluate the underlying immunosuppressive mechanisms in the TME. Existing in vivo models are ill-suited to answering the questions on mechanisms of cell migration and cell-cell interactions in the TME. Although in vivo models have the advantage of evaluating CAR T cell safety and efficacy, the models are largely immunosuppressed to allow for tumor engraftment and avoid rejection of human CAR T cells [18–22]. Furthermore, these models do not allow long-term investigation of the CAR T activity and locomotion in the solid tumor TME. Therefore, it is important to identify innovative models to evaluate CAR T cell function in the TME.

Advanced biomaterials [23–25] microfluidic [26–28] and bioprinting technologies [29,30] have contributed to our understanding of CAR T cells in solid tumors. Three dimensional (3D) in vitro models can recapitulate the solid tumor TME and allow reliable preclinical T cell investigation [31–33]. One 3D approach focuses on the tumor mass (e.g., tumor spheroid models) but not the physical and cellular surrounding [34–36]. T cells and tumor spheres are co-cultured in liquid media without an extracellular matrix (ECM) 3D support. Similar to conventional cell-monolayer 2D [37–42], these models do not provide the physiological spatial dimension and the physical barrier of the TME, known contributors to inefficient anti-tumor CAR T function in solid tumors [13]. Thus, the assays may not address the challenge of CAR T trafficking and could overestimate their efficacy. Another class of 3D models employs hydrogel platforms [27,43,44] where cells and tissues are stably supported in a continuum of extracellular matrix (e.g., synthetic or naturally derived). These platforms allow applications such as 3D migration [45], T cell cytotoxicity [46], or cancer differentiation [47]. However, they are limited to end-point analyses and have not demonstrated long-term investigation of the dynamic effector-target interaction.

CAR T cell therapies are designed to target surface antigens that are highly expressed on tumor cells compared with normal tissue to avoid the undesirable on-target off-tumor effect [48]. CD70 is the membrane-bound ligand of the CD27 receptor that is member of TNF receptor superfamily [49]. CD70 is a highly expressed marker in several solid tumors, including glioblastoma and osteosarcoma [50] Physiologically, CD70 is only upregulated by highly activated lymphocytes and dendritic cells (DCs), however, this expression is short-lived, rarely seen in cancer patients due to the heavy immunosuppressive state, and their depletion is not expected to induce overt immunodeficiency [51]. CD70 is also a marker associated with poor prognosis in number of solid tumors [52–54] These features make CD70 an attractive antigen for use in clinical trials that our group is pursuing (NCT05353530) [52,54].

In this study, we employed collagen-conjugated polyacrylamide liquid-like solid (COL1-LLS) [30,55], named in Vitro ImmunoTherapy Assays (iVITA), to characterize CAR T cell function in clinically relevant murine glioblastoma and osteosarcoma models. The COL1-LLS is an ensemble of microgels with interstitial space that act as microchannels for 3D cell migration. While each microgel particle presents a physical barrier toward the tumor site, the interstitial space forces the cells to explore paths of least resistance within the TME via chemotactic gradient [56]. Due to its optical transparency, the COL1-LLS allows long-term in situ visualization of cellular activities via real-time imaging. With iVITA platform, we can capture and quantify CAR T cell functions including dynamic trafficking, infiltration, activation, and killing of solid tumors. From these results, we derived a mathematical model demonstrating an inverse relation of CAR T infiltration and tumor elimination that can potentiate prediction of CAR T killing kinetics.

## MATERIALS AND METHODS

### CD70 expression and survival data of solid tumors

Primary patient gene expression data and matched normal tissue data as well as survival data were culled from the University of Alabama at Birmingham Cancer Data Analysis (UALCAN) that uses TCGA level 3 RNA-seq and clinical data from 31 different cancers [57,58]. The Human Protein Atlas RNA-seq data were used to provide information on survival of lung cancer and renal cancer [59–62].

### In Vitro Immunotherapy Assay

The LLS is an ensemble of transparent polyacrylamide microgels (Figure 1A). To facilitate cell adhesion and migration, we conjugated type I collagen on the surface of the LLS, namely COL1-LLS. For better visualization of the LLS architecture and presence of collagen on the microgel surface, a fluorescent anti-collagen antibody was used (Figure 1B). While the COL1-LLS supports single cells and tumor spheres in 3D (Figure 1C&D), the interstitial space between the particles provides random paths for T cell migration. We co-culture CAR T cells and tumor spheres in 3D in COL1-LLS to evaluate anti-tumor function via trafficking, expansion, and cytotoxic killing.

**Figure 1.**
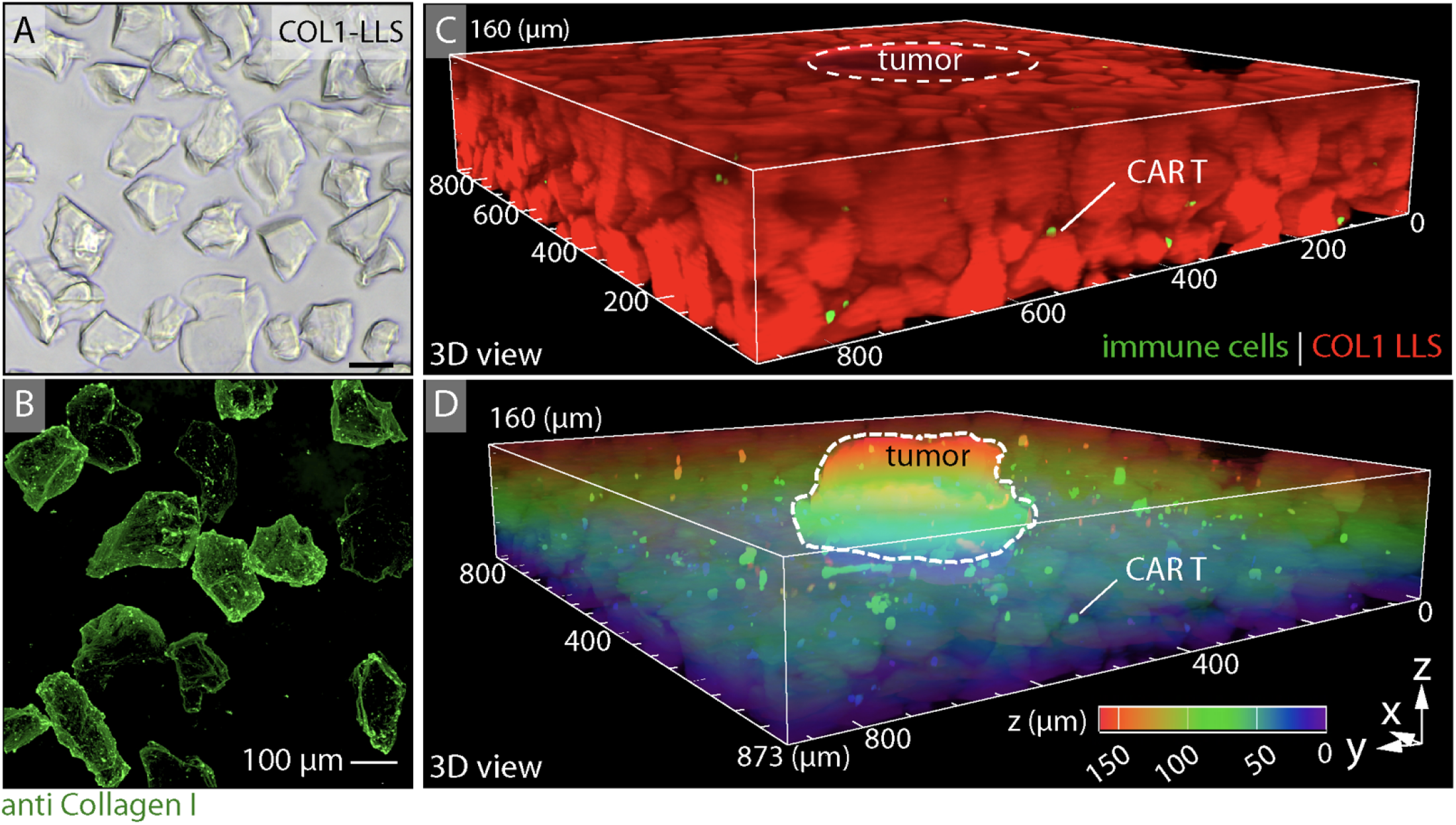
In Vitro Immunotherapy Assay (iVITA). A) brightfield image of type I collagen-conjugated LLS (COL1-LLS). B) Fluorescent image of COL1-LLS stained for anti-collagen antibody (green) confirms presence of the biomolecules on the surface of LLS. C) 3D view confocal image of iVITA platform showing dispersion of CAR T cells (green) around a tumor sphere in COL1-LLS (red). D) Depth coded max IP of (C).

### CAR Transduction of T Cells

The pMSGV8-mCAR construct contains the full CD27 sequence linked to the CD3z intracellular domain. A reporter gene, eGFP, is included in the construct to assess transduction efficiency and localization in vitro and in vivo assays. T cell transduction was done as previously described [63]. Briefly, murine T cells were isolated using magnetic selection according to manufacturer protocol (Pan T cell Isolation Kit II, Miltenyi, Germany) and activated with mouse anti-CD3 (5ug/ml) and mouse anti-CD28 (5ug/ml) (Gibco, Billings, MT, USA, 11132D) antibodies for 48-72 hours. Upon activation, T cells were transduced with pMSGV8-mCAR construct-containing plasmid, and CAR transduced T cells were cultured in recombinant human IL-2 (30U/ml)-containing complete T cell medium. Complete T cell medium contains RPMI (Gibco, Thermofisher, Waltham, MA), 10% FBS, 1% Non-essential amino acids (MEM NEAA, Gibco, Thermofisher, Waltham, MA), 1% penicillin/streptomycin (Gibco, Thermofisher, Waltham, MA), 1% sodium pyruvate (Gibco, Thermofisher, Waltham, MA), and 0.1% β-mercaptoethanol (BME, Gibco, Thermofisher, Waltham, MA). CAR T cells were ready for experiments after 48-72 hours of culture.

### Cell Culture

293T cell line (ATCC, CRL-3216) was used for lentiviral plasmid production encoding for mouse full length CD70 surface antigen (GenBank: U78091.1). This plasmid was used to transduce two murine solid tumor cell lines (KR158B and K7M2, ATCC). KR158B is a murine glioblastoma cell line. CD70-expressing KR158B (CD70^pos^ KR158B, kind gift of Dr. Jianping Huang) and wild-type (WT) KR158B cell lines were maintained in DMEM (Gibco, Thermofisher, Waltham, MA) without sodium pyruvate and 10% fetal bovine serum (FBS). K7M2 is a highly metastatic murine osteosarcoma cell line. CD70-expressing K7M2 (CD70^pos^ K7M2, kind gift of Dr. Elias Sayour) and wild-type (WT) K7M2 cell lines were maintained in DMEM with sodium pyruvate and 10% FBS. GP2-293 cell line (Takara, CA) was used as the viral packing cell line for plasmid encoding CAR construct. Cell cultures were maintained in a 5% CO_2_ incubator at 37°C for 2D and 3D experiments. All cell lines were pathogen-free according to IDEXX panel and approved by the internal EHS committee. The tumor spheres were formed by means of perfusion culture as previously reported [55]. In short, a solution of suspended single cells in inert LLS was plated into each well of a perfusion plate at a concentration of 10^6^ cells/mL. A medical suction device (i.e., a silicone bulb reservoir) was connected to create a “negative pressure” that gently perfused media through the interstitial space between LLS particles [55]. The continuous perfusion of media constantly supplied cells with nutrients while ensuring the proper removal of metabolic waste. Cancer cells aggregated by 48h and grew into tumor spheres ~400-600 μm in diameter by day 7.

### Co-culture Assay

For the 2D assay, CAR T cells were co-cultured with either wild-type or CD70-expressing tumor cells in a 1:1 ratio in 24-well plates without cytokines. CAR T cells were rechallenged with more tumor cells every three days until there was minimal evidence of CAR T cells identified by GFP signal to evaluate for exhaustion markers.

For 3D co-cultures, tumor spheroids were generated as previously described [55]. In short, suspended cancer cells (10^6^ cells/mL) were homogenously mixed in inert LLS and cultured in the Darcy plate by means of perfusion. The tumor spheroids were observed after 48h and ready by day 7. We co-culture CAR T cells with tumor spheres in 3D in type I collagen conjugated liquid-like solid (COL1-LLS) to evaluate anti-tumor function via trafficking, expansion, and cytotoxic killing. We homogenously mixed CAR T cells at a selected concentration in COL1-LLS and deposited the mixture into a glass-bottom 96-well plate. The tumor spheroids were then position into each well containing CAR T suspended in 3D in COL1-LLS. We then centrifuged the plate at gentle acceleration and deceleration setting. After centrifugation, we gently added growth media and secured the plate in a custom-made stage incubator on a Nikon A1R confocal microscope equipped with a high-definition Galvano scanner for long-term in situ imaging (~ 1-2 weeks). The CAR T and tumor cells can then be tracked, quantified, and analyzed. The anti-tumor activity from CAR T cells captured by this assay was also modeled mathematically to further predict CAR T-tumor interaction. At selected timepoints, we performed single-cell collection for immunophenotyping and collected supernatants for cytokine measurement. The co-cultures were maintained in incubators at 37°C and 5% CO_2_.

### Single cell collection and RNA sequencing

Single cell harvesting was performed using the CellCelector device (ALS, Germany). This technology utilizes immune fluorescence microscopy coupled with highly accurate single cell collection via capillary microfluidic system. Images are processed by a PC workstation. Individual cells were collected based on immunofluorescence and geographic position relative to tumor cells. The collected individual cells were placed in respective PCR tubes for subsequent downstream single cell RNA sequencing. Single-cell RNA library was prepared using QIAseq FX Single Cell RNA Library kit (Hilden, Germany) and carried out following manufacturer instructions. Transcriptome sequencing was conducted by Novogene Co., LTD (Beijing, China).

### Flow Cytometry

The following fluorochrome conjugated anti-mouse antibodies were used and included CD3 APC-Cy7 (BD Bioscience, New Jersey, USA), CD70 APC (Biolegend, San Diego, USA), PD1 APC (BioLegend, San Diego, USA), Tim3 APC (BioLegend, San Diego, USA). Flow staining was performed in FACS buffer (PBS + 2% FBS). For staining, 2 μL of antibodies (1μg/μL) were incubated with up to 10^6^ cells for 15-20min at room temperature. After washing samples with FACS buffer (x2), flow cytometry data were acquired using FACSCanto II (BD Bioscience) and analyzed using FlowJo Version 10.8. Given the clear separation of fluorochrome emission signals, compensation or fluorochrome-minus one (FMOs) were not used.

### Cytokine Measurement and ELISA Assays

Media supernatants were collected from in vitro co-cultures at 48 hours for evaluation of multiple cytokines, including inflammatory cytokines and chemokines using mouse IFNg DuoSet ELISA kit (R&D Systems, Minneapolis, MN, USA). In addition, Mouse Cytokine/Chemokine 44-Plex Discovery Assay^®^ Array (MD44) (Eve Technologies, Canada) was employed to measure 44 different cytokines/chemokines from effluent media.

### Microscopy

The in intro immune killing of tumor spheres were captured via long-term in situ imaging using a Nikon A1R HD25 confocal microscope equipped with a high-definition Galvano scanner. At selected timepoints, we performed single-cell collection for immunophenotyping and collected supernatants for cytokine measurement. The co-cultures were maintained in incubators at 37°C and 5% CO_2_. CAR T cells were manually segmented for cell tracking and then trained via Cellpose [64]. The identified CAR T cells were tracked via imageJ Linear Assignment Problem (LAP) tracker.

### Statistical Analysis

GraphPad Prism 9 software was used for statistical analysis. For comparison between the two groups, we used Student’s t-test. We used one-way and two-way ANOVA with multiple comparison tests for multiple-group comparisons. Statistical analysis was also included in the figure captions.

## RESULTS

### CD70 gene expression correlates with survival outcomes

Many solid cancers upregulate expression of the CD70 ligand compared with matched normal tissue cells, particularly esophageal carcinoma, cervical squamous cell carcinoma, head and neck squamous cell carcinoma, glioblastoma, sarcomas, and renal cancer (Figure 2A). In patients diagnosed with kidney cancer (n=877 patients; low expression= 185 vs high expression=792) and lung cancer (n=994 patients; low expression= 629 vs high expression= 365), it has been observed that high CD70 expression is associated with overall poorer survival (Figure 2B&C; p<0.001, respectively).

**Figure 2.**
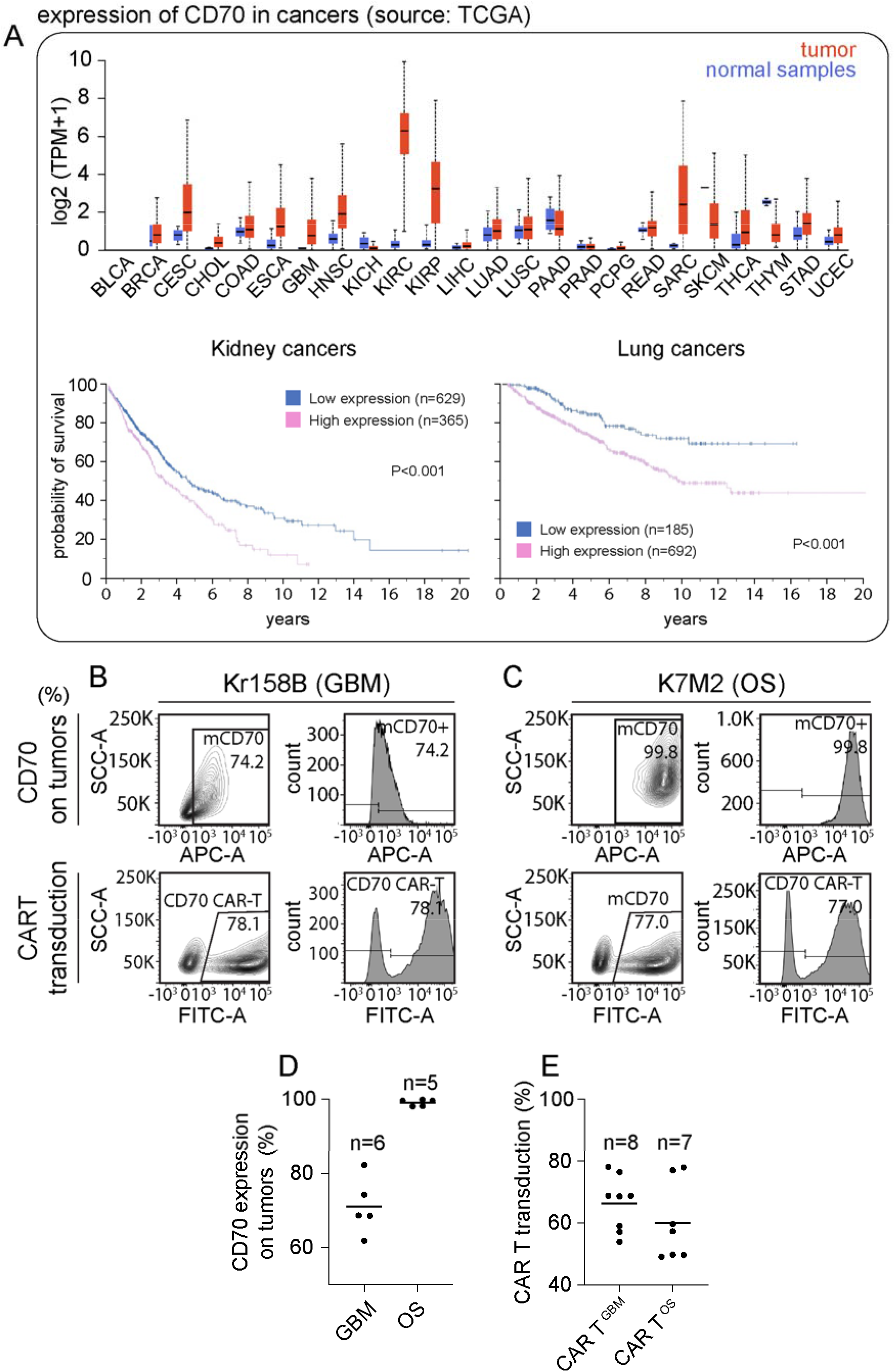
CD70 expression correlates with survival in patients with solid tumors. A) (top) CD70 gene expression for different solid tumors compared with normal matched tissues. Sources: The University of Alabama at Birmingham Cancer data analysis (UALCAN) and The Cancer Genome Atlas program (TCGA). (Bottom panels) survival rates of patients diagnosed with renal carcinoma and lung cancer with low and high CD70 expression. Source: The Human Protein Atlas program. B-C) Confirmation of CD70 (target) expression on cancer cells – B) glioblastoma (GBM) and C) osteosarcoma (OS) models - and (bottom) transduction of chimeric antigen receptor (CAR) construct in B) C57BL/6 mice - derived T cells (CAR T ^Kr158B^) and C) in Balb/c mice - derived T cells (CAR T ^K7M2^) as indicated by GFP reporter. On average, D) percentage of CD70 expression is 73% for glioblastoma and 99% for osteosarcoma models. E) Mean transduction of 66 % for CAR T ^Kr158B^ and 60% for CAR T ^K7M2^. The name of the cell lines and tumor models will be used interchangeably – GBM for Kr158B and OS for K7M2.

### Characterization of CD70^pos^ Expression and CAR Transduction Efficiency

Two murine solid tumor models were employed to demonstrate the reproducibility of tumor-immune interactions in iVITA. Murine glioblastoma (GBM) and osteosarcoma (OS) tumor spheres were created from KR158B and K7M2 cell lines, respectively. The CD70 expression of the target tumors was confirmed via flow cytometry prior to each experiment. The average target expression for GBM and OS tumors was 73% and 99%, respectively for all experiments (Figure 2B&C). For CAR T cell transduction, C57BL/6 mouse-derived T cells were obtained for the glioblastoma (GBM) tumor model denoted as CAR T^GBM^, and Balb/c mouse-derived T cells were used for osteosarcoma (OS) tumor model denoted by CAR T^OS^. The T cells from both models were transduced with a pMSGV8-mCAR construct containing a full-length CD27 sequence linked to the CD3z domain and eGFP reporter as previously described [52,54,63]. The transduction efficiency of the two CAR T backgrounds was assessed via flow cytometry and recorded as 66% for CAR T^GBM^ and 60% for CAR T^OS^ on average for all experiments (Figure 2D&E).

### CAR T Trafficking and Tumor Infiltration

During the first 14 hours of co-culture, CAR T cell migrate towards a tumor spheroid as captured via confocal time-lapse (Figure 3A). Tracking cell movement over time revealed higher CAR T cell velocity and infiltration when cocultured with their corresponding target tumors than the wild-type (WT) counterparts (Figure 3B-E). By means of confocal imaging, the projection of color-coded velocity tracks over time revealed the accumulation of CAR T cells inside a target tumor (Figure 3F). Quantification of time-lapse data confirmed the CAR T infiltration, as the percentage of total CAR T, was significantly higher for both CD70^pos^ GBM (35%) and CD70^pos^ OS (50%) tumors than their WT counterparts, at 15% and 21% for GBM and OS models, respectively (Figure 3C&E). In addition, evaluation of co-cultured supernatants marked upregulation of chemokines (CCL2, CCL3, CCL4, CXCL9, CXCL10) when CAR T cells encountered target tumors. These chemokines are known to be involved in T cell chemotactic trafficking and antitumor Th1 responses. Overall, chemokine secretion decreased over time for both tumor models except for CCL3 and CCL4 in the osteosarcoma tumor model (Figure 3G). In addition, we confirmed evidence of anti-tumor activity of CAR T cells via IFNγ which demonstrates remarkable difference in the first 72h from the co-culture with target tumors as compared to WT tumors for both models (Figure 3G).

**Figure 3:**
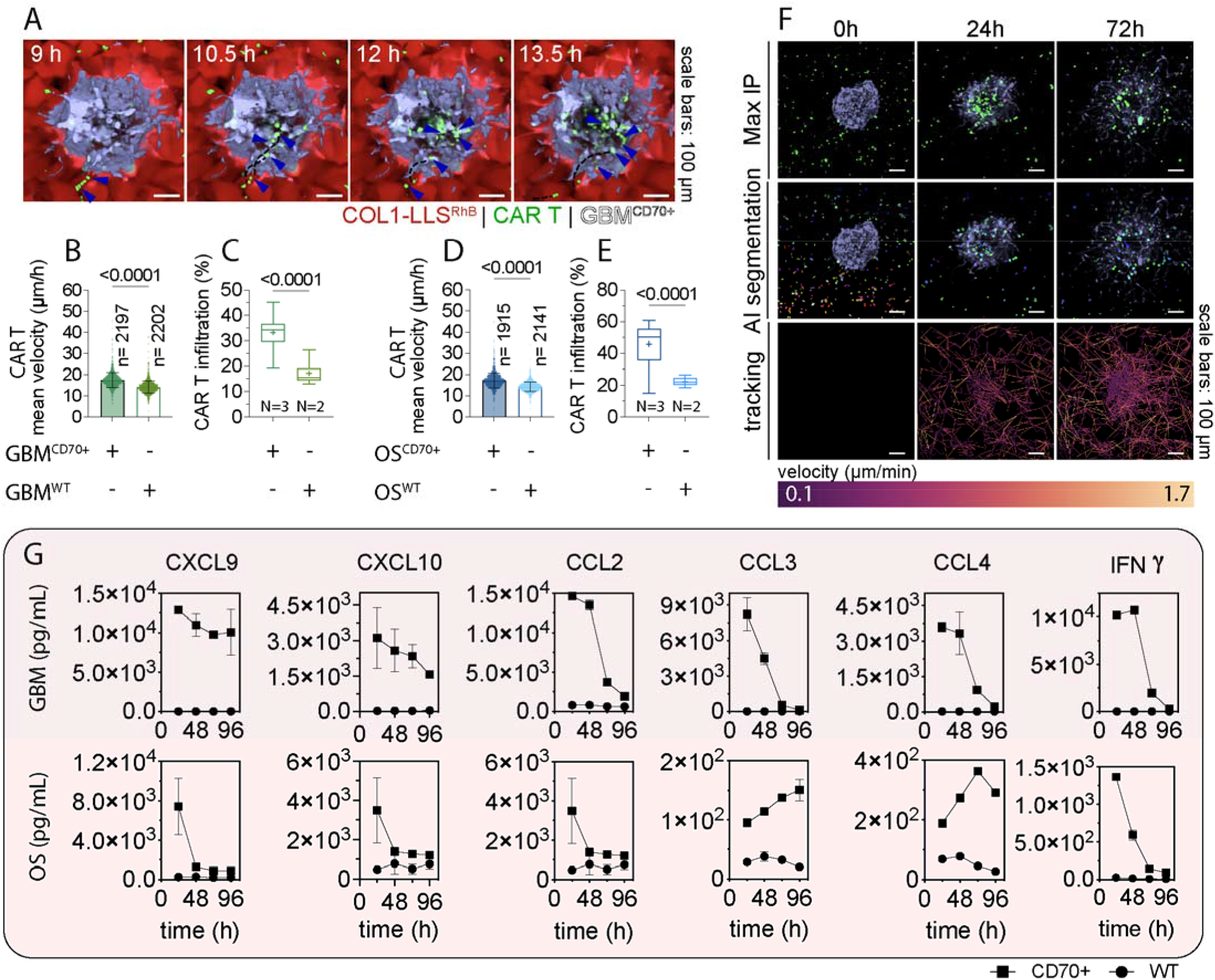
CAR T cell directed motion towards the target tumor. A) Representative confocal snapshots of CAR T cells infiltrating a GBM solid tumor in 3D. The panel shows CD70-specific CAR T cells (green) navigating through the supported COL1-LLS^RhB^ microgels (red) and infiltrating the target GBM tumor (white). Blue arrows indicate paths of CAR T migration. B, D) mean velocity of CAR T cells cocultured with CD70^pos^ tumors and the WT control for B) GBM and D) OS tumor models. The number of tracks (n) is indicated on the plots. The number of biological replicates is N=2. An unpaired two-tailed Student’s t-test was performed. Statistical significance with p values was indicated on the plots. C, E) Quantifying tumor-infiltrating CAR T cells on average from 0-72h as a percent of total CAR T cells for C) GBM and E) OS tumors. The box plots display 25^th^ and 75^th^ percentiles, a line at the median, and a plus sign at the mean, from the minimum to the maximum observation. Unpaired two-tailed Student’s t-test was performed (n=234, N=3, statistical significance with p values were indicated on the plots). F) Top panel: Maximum intensity z projection (top panel) showing snapshots of CAR T – GBM tumor interaction at 0, 24, and 72h. Middle panel: the segmentation of CAR T with colors indicated by individual CAR T cells at each frame. The segmentation employed a deep learning-based method (as discussed in the method section). Bottom panel: maximum intensity projection of the segmented CAR T cell velocity tracks overtime. The velocity gradient was color-coded, showing accumulation of CAR T inside the tumor. The segmented cells were tracked using Linear Assignment Problem (LAP) tracker at the maximum frame-to-frame linking and allowable track segment gap closing of 150 μm (~ 3 cell diameter). G) Evidence of chemotaxis and upregulation in migratory pathways for CAR T cells co-cultured with their target tumors for GBM (top panel) and OS (bottom panel). Noticeably, evidence of immune-mediated cytotoxic function is demonstrated via IFNγ detection.

### Indicators of CAR T Cell Expansion and Activation

We use CAR T expansion and interferon-gamma (IFNγ) secretion as indicators of activation and cytotoxicity. As shown in Figures 4A, CAR T cells in CD70^pos^ tumors were significantly higher than in WT tumors for both models. Next, we investigated the difference in CAR T expansion in the 3D COL1-LLS platform versus conventional 2D assays. In iVITA platform, the infiltrating CAR T cells constantly expanded up to 8-fold within the 72h period. This expansion, however, was not observed in WT tumors (Figure 4B-G). We defined 2D conditions as a cell monolayer cultured on a standard polystyrene or glass culture vessels submerged in liquid media. Flow cytometry evaluation revealed maximum CAR T expansion at 24h for GBM^CD70+^ (~3-fold) and 48h for OS^CD70+^ tumors (~4-fold) (Figure 4H&I). Furthermore, measurement of IFNγ secretion in both 2D and 3D conditions revealed a similar trend for activation and killing. In 2D co-culture, IFNγ levels reached a maximum of 5-6×10^3^ pg/mL at 24h and drastically decayed after 96h. In both co-culture conditions, while IFNγ levels from all samples decreased over time, those in 3D displayed a less drastic reduction immediately after the first 24h period (Figure 4J&K). Furthermore, T-cell clusters observed during expansion and activation suggest that this parameter can serve as an indicator for anti-tumor activity. Recently, studies have reported this phenomenon during T cell activation and killing of cancer cells [39,65,66]. In this study, CAR T cell clusters were observed during the co-culture with the target CD70^pos^ tumors but not the WT counterparts. The observation can be classified into two phases. During the first 24h, CAR T cells formed small clusters which rapidly increased, surrounded the tumor periphery, and formed a ring-like structure around the target tumors (Figure 4A). In the second phase (24-72h), the clusters converged and formed larger clusters (Figure 4A-E). These clusters negatively correlated with tumor size (Figure 4F&G) as the expansion of CAR T clusters and infiltration overwhelmed the tumor mass while the tumor noticeably reduced in size. In the WT tumors for both models, CAR T clusters were not detected, confirming the specificity of T-cell activation to only target CD70^pos^ tumors.

**Figure 4:**
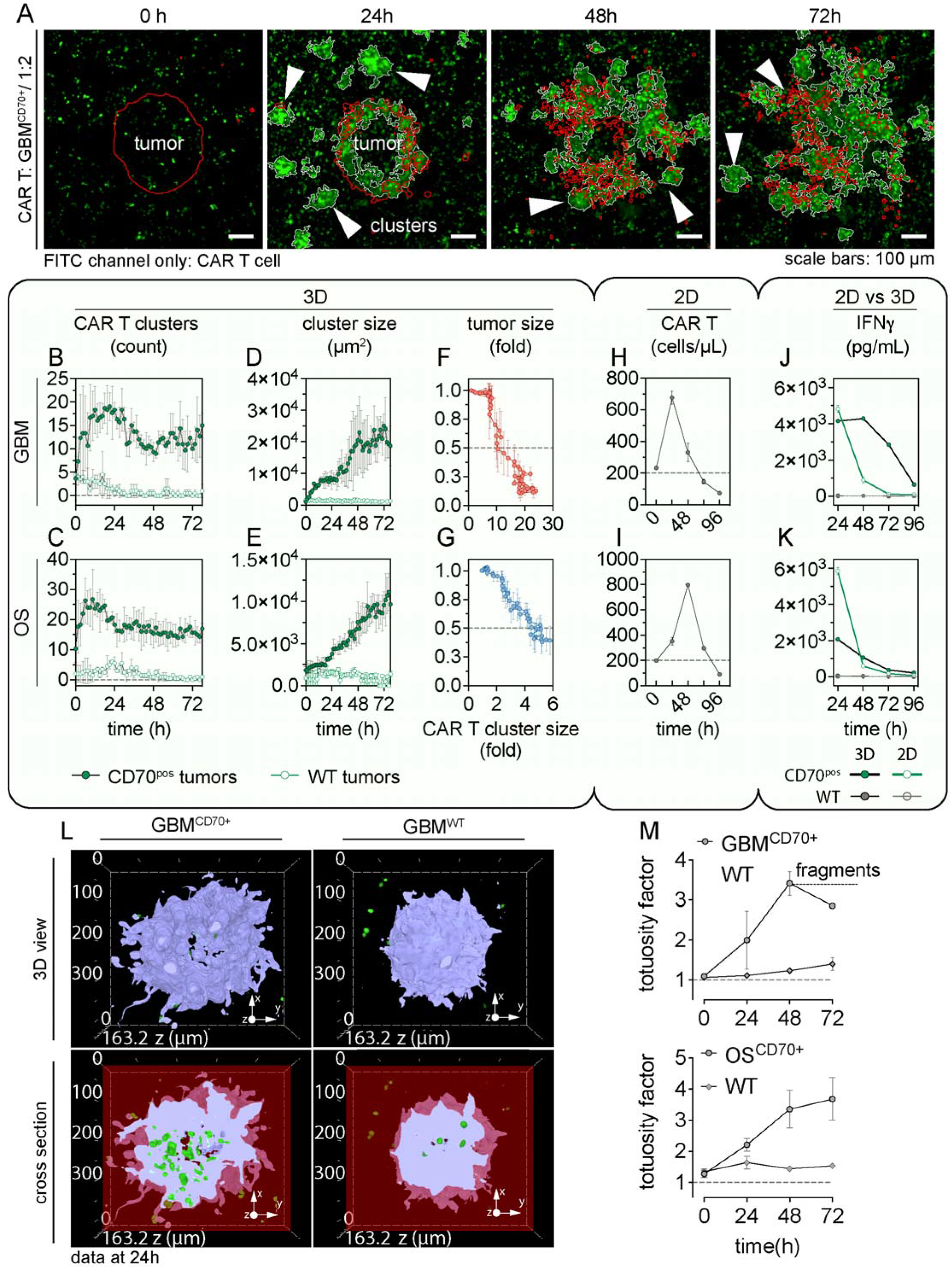
CAR T expansion, activation, and killing. A) Representative confocal timelapse images of the FITC channel (green) that show immune activation, expansion, and killing of the target tumors. The patterns of CAR T clustering and rapid expansion were observed in almost all conditions with efficient anti-tumor activity. White arrows locate CAR T cell clusters. B, C) The number of CAR T clusters rapidly increased during the first 24h and steadily maintained for more than 72h in B) GBM and C) OS models. D, E) Cluster size in all cocultures with target tumors steadily increased over time but not in the WT controls. F, G) Expansion of CAR T clusters revealed an inverse correlation with tumor size. (n=3 for all CD70^pos^ samples and n=2 for all WT samples). H, I) Endpoint flow cytometry data measuring CAR T expansion after 96h of CAR T – cancer cells coculture in the 2D assay for both tumor models. J, K) A comparison in IFNγ secretion (pg/mL) for 2D vs 3D for J) GBM and K) OS. The number of biological replicates is N=2. L) Top row: confocal 3D snapshots of GBM tumors after co-cultured with CAR T cells for 24h showing highly tortuous tumor margin of CD70^pos^ tumor as compared to the WT counterpart. Bottom row: cross-section view (z depth: 70 μm) of the sample on the top row exposing infiltrating CAR T cells within the tumor mass. M) Measurement of tumor tortuosity factors revealed more than 3-fold change for the CD70^pos^ tumors within the first 48h of coculture. The tumor tortuosity factor was calculated as a ratio between the perimeter of the tumor outline to the perimeter of a circle of the same pixel area. Data was obtained from GBM samples (n=3) at an initial E:T ratio of 1:2, and from osteosarcoma (K7M2) samples (n=3) at an initial E:T ratio of 1:1. Biological samples for each group (n= 3), and technical repetitions (n= 3) were performed.

An interesting observation from our study was the tortuous morphology that the target tumors adopted upon specific engagement with CAR T cells (Figure 4L). Upon an immune attack, cancer cells from CD70^pos^ tumors from both GBM and OS models migrate away from the tumor mass resembling an accelerated tumor invasion process. For CD70^pos^ tumors from both models, the cancer cells at the tumor periphery individually and collectively migrated away from the tumor mass upon CAR T infiltration, resulting in a tortuous tumor morphology distinctly different from all WT tumors. We defined the tumor morphology by a tortuosity factor that is the ratio of the tumor perimeter and the perimeter of an equivalent circle of the same tumor area. This factor was consistently 3 to 4-fold higher for the CD70^pos^ tumors than WT tumors of both models when cocultured with CAR T cells (Figure 4M). The 3D cross-section views revealed a noticeable amount of immune infiltration in the CD70^pos^ tumors relative to the WT tumors (Figure 4L) which remained intact with insignificant CAR T infiltration and invading cancer cells.

### Tumor Elimination and Tortuosity Directly Correlated with CAR T Infiltration

From our experiments with 2D assays for both tumor models, we seeded single CAR T cells homogenously into the iVITA assay such that the effectors to target ratio (E:T) was 1:2 for GBM and 1:1 for OS tumor. As shown in Figures 5A&B, CAR T cells and tumor spheroids were digitally segmented by their respective fluorescent markers (Figure 5, bottom rows of each panel). Figure 5A&B, top panels, shows qualitative evidence of the immune-mediated killing of target (CD70^pos^) tumors for both GBM and OS models. CAR T cells rapidly formed clusters during the first 5 hours, localized at peripheral regions of the target tumor, and grew more prominent within a 24h period. From 24-72h, CAR T cells, as single cells and clusters, trafficked to the periphery of the target tumors, combined, and carried out tumor elimination process leading to a significant reduction in tumor size. Such activities were not observed for CAR T cells in WT tumors (Figure 5A&B, bottom panels) validating the specificity and efficacy of CAR T killing of solid tumors in iVITA.

**Figure 5.**
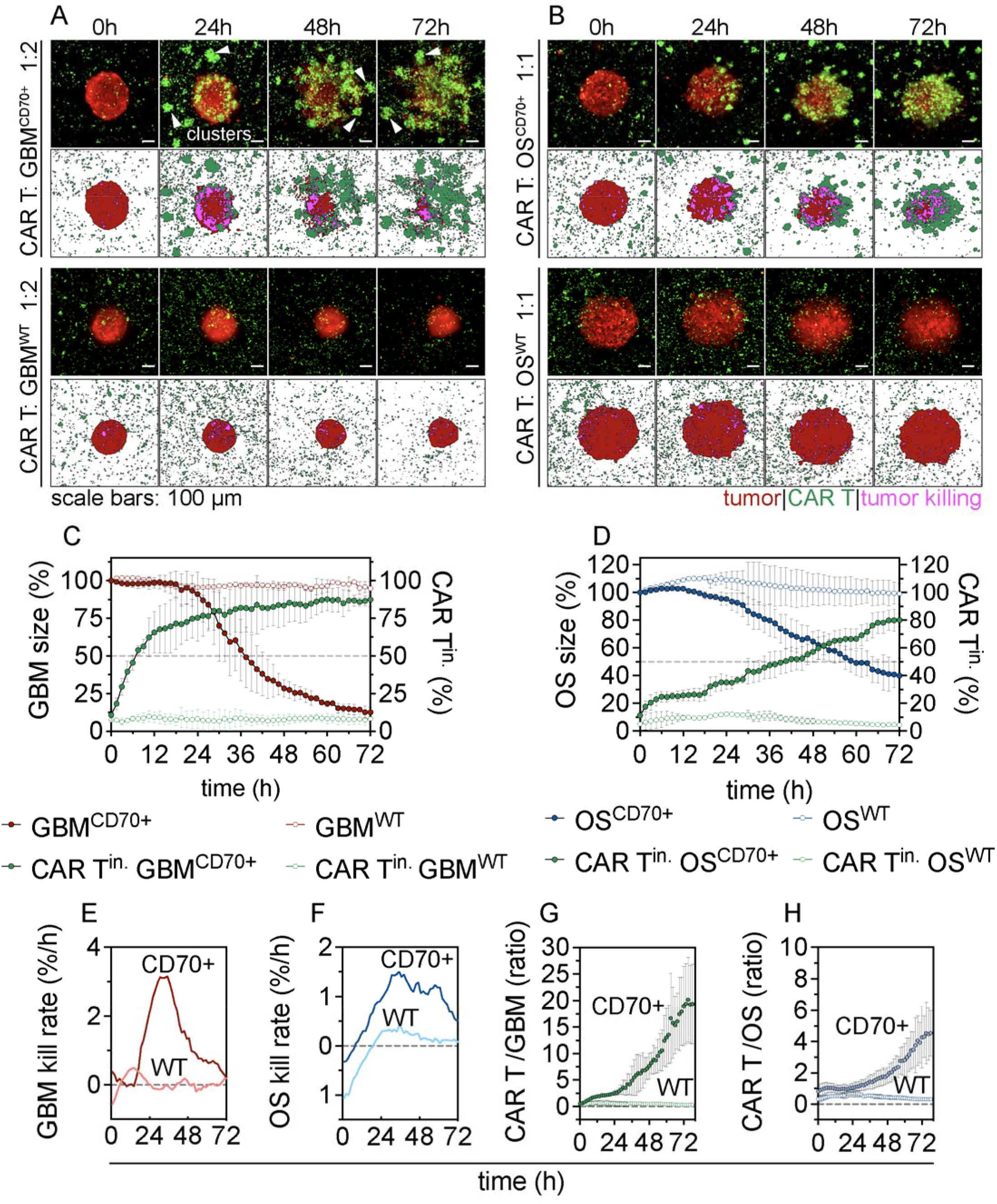
Quantification of CAR T-tumor interactions. Top rows: confocal timelapse images at 0, 24, 48, and 72h showing dynamic immune-cancer interactions and antitumor activities of CAR T cells in A) GBM and B) OS tumor models. The bottom row of each group represents a digital reconstruction of the confocal data with distinct segmentation of tumor and immune cell populations. The initial CAR T to tumor cells ratio or effector to target (E:T) ratio is indicated on the graph. C, D) Quantification of CAR T–tumor interactions showing inverse correlation between tumor size and infiltrating CAR T for C) GBM and D) OS tumors. The percentage of tumor size (left axis) and CAR T cell in the tumor (right axis) was normalized to the original tumor size at time 0. E, F) Killing rates over time were calculated as derivatives from C and D, respectively. G, H) The measured CAR T to cancer cells ratios (E:T) as a function of time. The E:T ratios dynamically changed over time. In the case of WT tumors, the ratios remain constant.

Quantifying tumor-immune interaction revealed an inverse correlation between tumor size and CAR T infiltration (Figure 5C&D). As the infiltrating CAR T cells increased, the tumor size decreased over time for both GBM and OS models. Noticeably, CAR T infiltration in the GBM model increased rapidly for the first 24 hours compared to the OS model. This was reflected in the reduction rate of tumor size, defined as tumor killing rate, which peaked at a maximum of ~3% of original tumor mass per hour (%/h^-1^) within 24-48h (Figure 5C&D). On the other hand, the killing rate for OS tumors was lower (~1.5 %/h) and remained relatively consistent for an extended period from 24-60h (Figure 5E&F). For both models, WT tumors did not change in size and the CAR T cell did not increase confirming the trivial (background noise) tumor-killing rate throughout the experiment (Figure 5E&F). Furthermore, the ratio of CAR T to cancer (areas) exponentially increased over time (Figure 5G&H), resulting from the noticeable expansion of CAR T cells and shrinkage of target tumors.

### Feasibility of identifying CAR T cell gene pathways

We demonstrated the feasibility of single cell collection that allows for evaluation of CAR T cell gene profiles relative to their role and location in the co-culture. The capability of the technology used to harvest a single cell in a culture is shown in Figure 6A. A device developed by our group with similar capabilities will be evaluated in future studies of tumor and CAR T cell biology (Figure 6A). We collected CAR T cells that were infiltrating and actively killing tumor cells -defined as “proximal” and CAR T cells that were away from the target tumors - defined as “distant”. We evaluated three pathways including T cell exhaustion, T cell cytotoxicity, and chemokine signaling. Even though the number of samples tested was limited to make any definitive conclusion, we observed a genetic profile that provides a differential signature to better understand the biology of CAR T cell relative to their activity in a 3D co-culture. There was increased gene expression of chemokine genes including CCL2, CXCL9, CXCR4, CCL20, CCR9 and Jak2 genes in CAR T cells that were co-cultured with CD70^pos^ tumors. These genes are known to be associated with T cell trafficking and infiltration of tumors [58,67–69]. Genes involved in cytotoxicity pathways were upregulated in CAR T cells co-cultured with CD70^pos^ tumors. The upregulation included granzyme and Fas-related genes. We also observed higher expression of exhaustion genes in the CAR T cells cultured with wild type tumors (CD70^neg^) compared with conditions where CAR T cells engaged with CD70^pos^ tumors. When we compared proximal-CAR T cells with distant-CAR T cells, more chemokine signaling- and cytotoxicity-related genes were upregulated. However, no marked differences were identified when T cell exhaustion genes were analyzed (Figure 6B).

**Figure 6:**
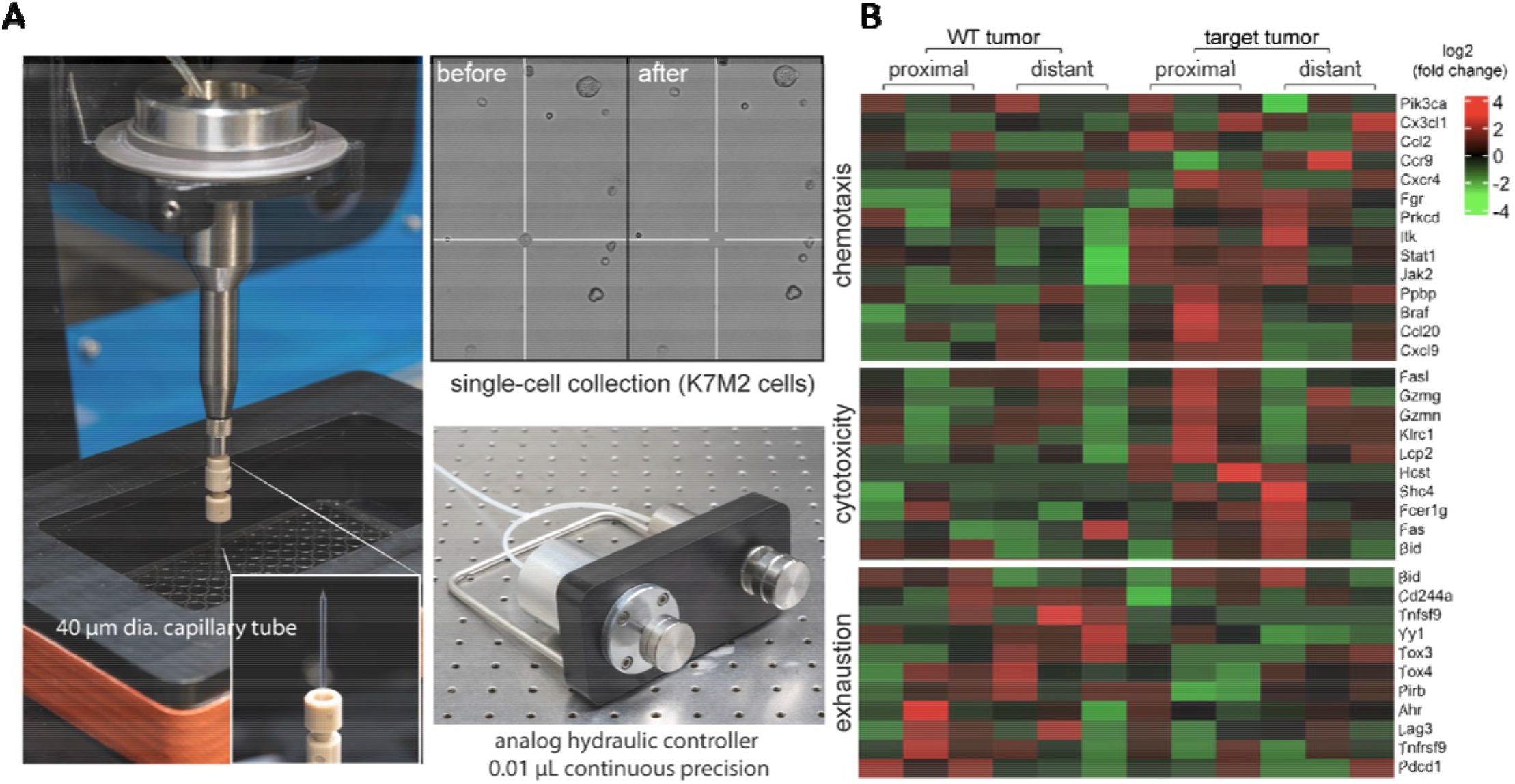
A) Single-cell collection and transcriptomic profiling of CAR T cells. During CAR T – tumor co-culture experiment, we collected single CAR T cells (1 cell per sample) at two distinct spatial locations relative to the tumor mass. We define those far away from the tumor (> 500 μm) “distant” CAR T cells and those at the vicinity (at the edge or on the tumor) “proximal” CAR T cells. B) We collected N=3 single cells per condition. We then performed transcriptomic profiling corresponding to three T cell pathways including chemokine signaling, T cell cytotoxicity, and T cell exhaustion. Values are represented in log-fold expression.

### Mathematical Modeling of CAR T-Mediated Elimination of Solid Tumors

In this model, CAR T cells are assumed to be effector T cells undergoing activation and expansion. An exponential function can then describe the number of infiltrating CAR T cells (N_i(t)_).

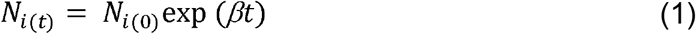

where N_i(0)_ represents the number of infiltrating CAR T cells at time t= 0, *β* is the rate of CAR T proliferation/recruitment to the tumors [70]. CAR T cell exhaustion is not included. On the other hand, the number of cancer cells (N_c(t)_) is dependent on the natural proliferation/apoptosis and cell death caused by CAR T-mediated killing. We arrive at the following expression for cancer cells, Equation 6-2.

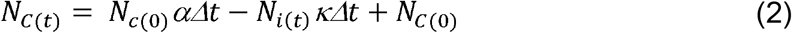

In the above expression, *α* presents the cancer proliferation rate based on experimental data and literature [71,72], and *κ* the immune killing rate[70–72]. Therefore, the rate of change of cancer cells can then be expressed as:

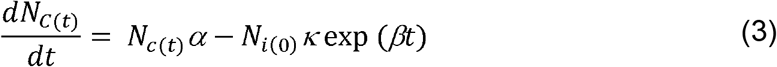

Based on the boundary condition assumption at time t=0, *N*_*c*(*t*)_ = *N*_*c*(*0*)_, the final form of a general solution, Equation 6-4, provides a mathematical model of the evolution of the tumor upon an immune attack.

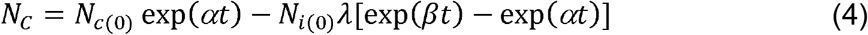

where 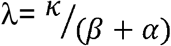 shows the relationship between the kill rate, cancer proliferation rate, and CAR T cell proliferation rate. Figure 8 shows that the model is conservatively predictive and demonstrates strong agreements with experimental data.

**Figure 7.**
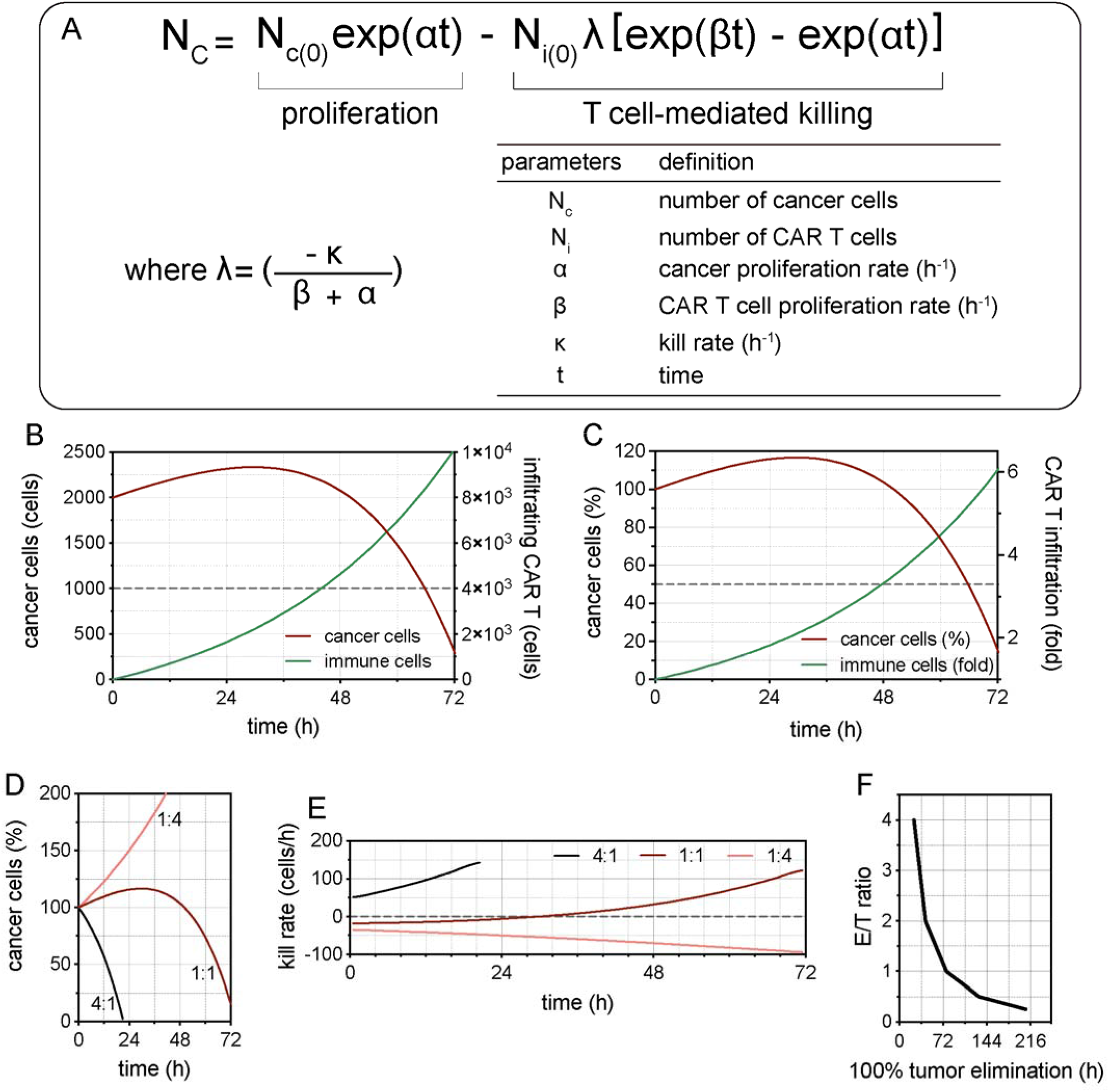
Mathematical model of in vitro tumor-immune interaction. A) Mathematical description of cancer cell number in response to T cell killing. The dynamic profile of the tumor can be determined by proliferation and T cell-mediated killing. B) Theoretical prediction of the inverse correlation between cancer cells and infiltrating CAR T cells. C) The same relation is described in terms of the percent of the original tumor mass and CAR T fold expansion. D) The dynamic profile of cancer cells, as a percentage of the original, as a function of time at different E:T seeding ratios. E) Rate of tumor killing was calculated as the first derivative of D). F) Theoretical prediction of complete tumor elimination at various E:T ratios as a function of time (h). Note: The model does not account for T cell exhaustion and natural apoptosis from both immune and cancer cell populations.

**Figure 8.**
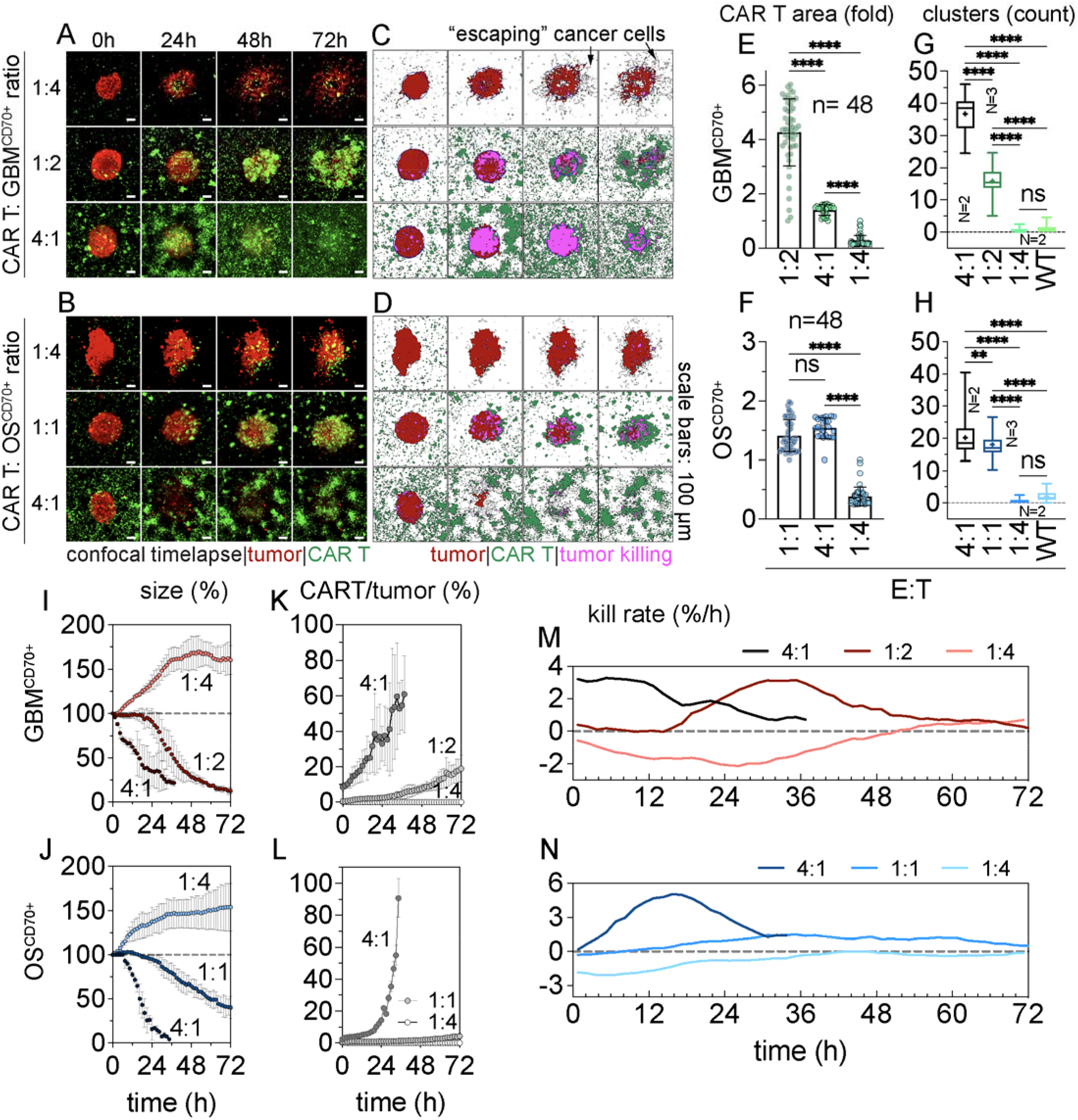
Sensitivity of anti-tumor activity to various CAR T: cancer cell (E:T) ratios. A,B) confocal time-lapse images of CAR T – tumor co-culture in iVITA at different E:T ratios. C,D) show the digital image reconstruction of confocal data quantifying bulk tumor mass, migrating single cancer cells, immune cells, and immune cell clusters, and killing activity over time. A,C) GBM CD70^pos^ tumors and B,D) OS CD70^pos^ tumors were co-cultured with their respective CAR T cells at different concentrations corresponding to E:T of 1:4, 1:2, and 4:1 for the GBM model and E:T of 1:4, 1:1, and 4:1 for the OS model. At 72h and E:T= 4:1, almost 100% tumor elimination was observed in both models. E-H) CAR T expansion as a function of initial E:T seeding ratios. E,F) CAR T expansion on average in both models from 0-72h. For each seeding E:T, CAR T expansion at each time point was normalized to the CAR T at time 0, and the average was calculated for all frames. G,H) the number of CAR T clusters counted every 1.5h for 72h for each group. The box plots display 25th and 75th percentiles, a line at the median a plus sign at the mean, from the minimum to the maximum observation. I,J) Quantification of tumor size over time at the initial E:T= 1:4, 1:2, and 4:1 for I) GBM model and at E:T= 1:4, 1:1, and 4:1 for J) OS model. K,L) The E:T ratio dynamically changed over time. The CAR T expansion and tumor-killing were presented by the exponential increase of E:T ratios. M,N) tumor killing rates were calculated as derivatives from I) and J), respectively. Statistical analysis was performed using Ordinary One-Way ANOVA. (n=3 unless indicated otherwise, ** = p < 0.01, and **** = p < 0.0001).

In addition, the model also predicts complete tumor elimination as a function of effector-to-target ratios (E/T) (Figure 7). For E/T ratios of 4:1 and 1:1, the model predicted the time to complete elimination as ~36h and 72h, respectively (Figure 7D-F), resembling the aforementioned data. The prediction, however, became less accurate at low E/T. The model theoretically indicated a possible tumor elimination after five days of coculture for an E/T ratio at 1:4. However at such a ratio, immune exhaustion became evident in experimental data where there was minimal detection of immune infiltration, expansion, and killing, and tumor size did not show sign of regression.

### Evaluation of Anti-tumor Function In Response To E:T Seeding Ratios

While CAR T anti-tumor function was evident for both models, we asked whether a complete tumor elimination can be observed. iVITA assays were set up to evaluate the sensitivity of anti-tumor activity in response to different E:T seeding ratios and the possibility of a complete tumor elimination. The CD70^pos^ tumor spheres (~10^3^ cells) from the two murine GBM and OS solid tumor models were positioned in COL1-LLS where CAR T cells were uniformly distributed in 3D. Three E:T ratios at 1:4, 1:2, and 4:1 were selected for the GBM and the ratios at 1:4, 1:1, and 4:1 for the OS (Figure 8A&B). Over time, qualitative data of cancer-immune interaction revealed two observable modes of CAR T infiltration and killing: inside-out and outside-in attacks. Single CAR T cells that successfully infiltrated the tumor mass proliferated and carried out the anti-tumor activity from the inside of the tumors (Figure 8A&B, first rows). On the other hand, CAR T cells at higher seeding ratio (E:T of ≥ 1:2) also contributed to an outside-in attack. Similar to previous experiments, it was observed that CAR T cells trafficked towards a target tumor both individually and as clusters (Figure 8A&B, middle and bottom rows). The clusters then combined to form larger ones at peripheral regions of the tumor mass and rapidly overwhelmed the target tumor. As shown in Figures 8C&D (right panels-middle and bottom rows), the overlapping regions of CAR T cells and the tumor (segmented in magenta and defined as tumor killing zone) demonstrated a ring-like pattern around the tumor periphery and immune infiltration increased over time. The outside-in attack was the dominant mode of tumor elimination for E:T≥1:2. Within 72 hours of co-culture, CAR T cells overwhelmed the tumor mass, and a complete tumor elimination was observed in the 4:1 conditions for OS models. At a low E:T ratio (1:4), CAR T cells trafficked to the tumors as single cells, and there was minimum cluster formation in both CD70^pos^ and WT tumors.

The total CAR T expansion was at maximum 4-fold and 1.5-fold for GBM (at 1:2) and OS (at 4:1) tumors, respectively (Figure 8E&F) for the first 48h period. The number of CAR T clusters from both CD70^pos^ tumor models were relatively high, > 20 for both tumor models at E:T ≥ 1:2, respectively, compared to low E:T seeding ratio (1:4) and WT tumors of both models (Figure 6G&H). At E:T= 4:1, CAR T clusters were observed immediately within the first hour of the co-culture. The clusters grew larger over time, consistent with previous observations (Figure 5). Due to T cell expansion and tumor elimination, the E:T ratios dynamically changed over time and showed an exponential increase, demonstrating significant sensitivity of this parameter to the anti-tumor function (Figure 8I&J).

Quantification of tumor killing at different E:T ratios revealed dynamic tumor elimination over time. CAR T cells at E:T ratio ≥ 1:2 for GBM and ≥ 1:1 for OS demonstrated efficient trafficking, expansion, infiltration, and killing of target cancer cells (Figure 8K&L). On the other hand, at a low E:T seeding ratio (1:4), antitumor activities of CAR T cells were short-lived and did not demonstrate evidence of tumor regression. At an intermediate ratio (1:2 for GBM and 1:1 for OS), the tumor size regressed after 24h of the coculture, confirming the two phases of an immune attack. The first phase comprised CAR T trafficking, infiltration, and expansion (Figure 8A,B,K,L). The second phase revealed rapid tumor eliminations (Figure 8A,B,I,J,M,N). The tumor-killing rate reached a maximum at 24 and 48h in GBM (1:2) and OS (1:1), respectively. At E:T= 4:1, tumor elimination happened almost immediately, suggesting tumor elimination was sensitive when E: T was sufficiently high. The study indicated that efficient anti-tumor function was highly dependent on the E:T ratios, which could serve as helpful information for predicting CAR T-cell doses needed to treat patients with advanced solid tumors.

## DISCUSSION

Despite the success of CAR T cells treating patients with hematological malignancies, CAR T technologies have only provided modest to no survival benefit to patients with solid tumors largely due to the heavily immunosuppressive TME [13,73]. A critical component to better understand these obstacles lays in the development of novel ex vivo technologies that can allow reliable study of cancer and immune cell dynamic behaviors. In this study, we hypothesized that cancer and CAR T cell interactions can be qualitatively and quantitively characterized ex vivo at the single cell resolution using a 3D microgel technology. Our bioconjugated liquid-like solid microgels demonstrated high resolution evaluation of the in vitro antitumor activity of CAR T cells in an environment that replicates the physical barriers imposed by solid tumors.

A major challenge of CAR T cell therapy for solid tumors comes from inefficient infiltration into the tumor mass. In the TME, the dense, fibrous, and crosslinked ECM proteins significantly contribute to the barriers hindering CAR T cell activity [74]. In physiological conditions, CAR T cells must efficiently overcome inherent barriers of the solid tumor TME to carry out anti-tumor function [13]. In iVITA we reconstructed ECM components as building blocks of protein-conjugated LLS (e.g., COL1-LLS) to investigate T cell locomotion in the tumor spheroid. The randomly tortuous microchannels formed by the interstitial space of COL1-LLS enable cell migration independent of proteolytic capability while the microgels impose physical barriers that CAR T cells must overcome to reach the target tumor. Naturally, this mode of locomotion, known as amoeboid, employs migration through paths of least resistance, actomyosin contractility, coordinated membrane extrusion, and contact-induced traction forces on their surroundings to squeeze and navigate through accessible pores [75–77]. Furthermore, the immune cells must be able to sense the chemotactic gradient and traverse through complex and narrow interstitial space networks to meet their target cancer cells [13]. Upon antigen specific immune attack, the observed increased secretion of chemokines involved in CCR2, CCR5 and CXCR3 pathways [68,78–80] in the target tumor conditions further support a chemotaxis-mediated CAR T migration to the tumor. These findings raise the questions to be confirmed in future studies of which chemokine pathway is critical for CAR T cell trafficking within the solid TME.

We observed CAR T cell cluster formation that directly correlated with antitumor activity specific to the tumor target which was not observed in negative surface antigen tumors. These CAR T cell clusters might resemble tertiary lymphoid structures (TLSs) which have been reported as positive prognostic factors associated with survival outcomes [81,82]. The TLSs could be well organized, lymph node-like structure with various cell types or simply lymphocyte aggregates [83]. Despite being suggested as a positive predictor of patient survival, the fundamental anti-tumor mechanism of TLSs has not been clearly explained. Therefore, CAR T cell clusters in our system could potentially serve as a reliable indicator of CAR T activation and antitumor function.

An immune attack could cause an abundant cytokine and chemokine release leading to a stress-associated tumor progression and invasion [84]. The chaotic and stressful T-cell-mediated cytotoxicity against the target tumors could also disrupt the cell-cell interaction, thereby promoting tumor dissemination. In iVITA, cancer cells individually and collectively migrated out of tumor mass upon CAR T cell attack which suggests an immune evasion mechanism. The cancer cell local migration led to a tortuous tumor mass only observed in the presence of antigen specific CAR T killing. These findings may provide a novel understanding of the relationship between cancer-immune interaction, immune escape, and cancer progression [85,86].

With the qualitative and quantitative data from iVITA, we developed a mathematical equation to predict antitumor function of CAR T cells against solid tumors. The mathematical model showed reasonable predictions and potential application to different solid tumor models. However, this model provides an ideal mathematical interpretation, and therefore is not without limitation. The model assumes infinite cell proliferation capability with trivial T cell exhaustion. Due to the overwhelming CAR T mediated killing and observable tumor reduction in a relatively short timeline, we did not consider immune exhaustion a governing variable in our equation. Recently, a human clinical trial demonstrated that CAR T cells dose positively correlates with long term survival benefits [87]. Thus, this mathematical description might inform clinical trials on predicted cell dose for optimal antitumor killing.

The iVITA platform can be coupled with a single cell collection technology for further immune and gene profiling to help identify the CAR T cells with optimal antitumor function. At the genomic level, our results show higher cytotoxic and trafficking chemokine gene upregulated for CAR T cells that encountered their target tumor. Consistent with our mathematical assumptions, we observed lower exhaustion gene signature for CAR T cultured with antigen positive tumors compared with conditions treating antigen negative tumors. The CD70-CD27 axis is an activating pathway and hence, the lower expression of exhaustion genes in CAR T cells cultured with CD70^pos^ tumors was expected [88]. Taken together, this platform could guide the design of a CAR T cell with killing, trafficking, and persistent characteristics capable of overcoming the immunosuppressive TME of solid tumors.

## CONCLUSION

Thus far, conventional models and current biomarkers have not been able to predict treatment outcomes. A more physiological ex vivo system including various cell subsets and microenviromental components will potentiate the faithful reproduction of the tumor immune interactions happening in cancer patients. The presented in vitro model enables mechanistic studies of CAR T cell trafficking and function in solid tumors. In addition to the patient-derived xenografts, our system represents a promising alternative to investigate the solid tumor TME with the advantage of being a more affordable technology. This model may allow in vitro assessment of autologous CAR T cells against patient-derived tumor explants potentiating advancement in personalized CAR T cell design and production. The advantages offered by three-dimensional cell culture technologies might position them as reliable co-clinical trial correlates to evaluate therapeutic response and toxicity.

## ACKNOWLEDGMENTS

We thank you the funding support from the National Science Foundation Graduate Research Fellowship (DGE-1842473, D.T.N.), Merck & Co. (W.G.S. and D.T.N.) NCI R37CA251978 (E.J.S.), Rally Foundation (P0216283, E.J.S.), and a Bankhead Coley Research Grant (P0172137, E.J.S.), FDOH Live Like Bella Discovery grant (22L07, P.C.), UFHCC / CTSI KL2TR001429 (P.C.) and UL1TR001427, and the Stop Children’s Cancer foundation (F023891, P.C.), Department of Defense Cancer Research Program (W81XWH-20-1-0726, J.H.)

## AUTHOR CONTRIBUTIONS

DTN & RL: Conceptualization, methodology, validation, formal analysis, investigation, data curation, writing – original draft, writing – review & editing. EOR, AP: Conceptualization, methodology, validation, formal analysis, investigation, data curation. DP: formal analysis, investigation, data curation. SQ: investigation, data curation. NTYN, JML, GRG, RAS, LJ, HT, AW, SP: investigation. JL: writing – review & editing. DAM, EJS: conceptualization, methodology, supervision, funding acquisition, writing. JH, PC, WGS: Conceptualization, methodology, validation, formal analysis, investigation, data curation, writing – original draft, writing – review & editing, supervision, funding acquisition.

## REFERENCES

1. Siegel, R.L.; Miller, K.D.; Fuchs, H.E.; Jemal, A. Cancer statistics, 2022. CA. Cancer J. Clin. 2022, 72, 7–33. 10.3322/caac.21708.

2. Costello, J.C.; Heiser, L.M.; Georgii, E.; Gönen, M.; Menden, M.P.; Wang, N.J.; Bansal, M.; Ammad-Ud-Din, M.; Hintsanen, P.; Khan, S.A.; et al. A community effort to assess and improve drug sensitivity prediction algorithms. Nat. Biotechnol. 2014, 32, 1202–1212. 10.1038/nbt.2877.

3. Mitchell, D.A.; Batich, K.A.; Gunn, M.D.; Huang, M.N.; Sanchez-Perez, L.; Nair, S.K.; Congdon, K.L.; Reap, E.A.; Archer, G.E.; Desjardins, A.; et al. Tetanus toxoid and CCL3 improve dendritic cell vaccines in mice and glioblastoma patients. Nature 2015, 519, 366–369. 10.1038/nature14320.

4. Ogando-Rivas, E.; Castillo, P.; Jones, N.; Trivedi, V.; Drake, J.; Dechkovskaia, A.; Candelario, K.M.; Yang, C.; Mitchell, D.A. Effects of immune checkpoint blockade on antigen-specific CD8+ T cells for use in adoptive cellular therapy. Microbiol. Immunol. 2022, 66, 201–211. 10.1111/1348-0421.12967.

5. O’Leary, M.C.; Lu, X.; Huang, Y.; Lin, X.; Mahmood, I.; Przepiorka, D.; Gavin, D.; Lee, S.; Liu, K.; George, B.; et al. FDA Approval summary: Tisagenlecleucel for treatment of patients with relapsed or refractory b-cell precursor acute lymphoblastic leukemia. Clin. Cancer Res. 2019, 25, 1142–1146. 10.1158/1078-0432.CCR-18-2035.

6. Bouchkouj, N.; Kasamon, Y.L.; Claro, R.A. de; George, B.; Lin, X.; Lee, S.; Blumenthal, G.M.; Bryan, W.; McKee, A.E.; Pazdur, R. FDA approval summary: Axicabtagene Ciloleucel for Relapsed or Refractory Large B-cell Lymphoma. Clin. Cancer Res. 2019, 25, 1702–1708. 10.1158/1078-0432.CCR-18-2743.

7. Wang, M.; Munoz, J.; Goy, A.; Locke, F.L.; Jacobson, C.A.; Hill, B.T.; Timmerman, J.M.; Holmes, H.; Jaglowski, S.; Flinn, I.W.; et al. Three-Year Follow-Up of KTE-X19 in Patients With Relapsed/Refractory Mantle Cell Lymphoma, Including High-Risk Subgroups, in the ZUMA-2 Study. J. Clin. Oncol. 2022, 94. 10.1200/JCO.21.02370.

8. Abramson, J.S.; Palomba, M.L.; Gordon, L.I.; Lunning, M.A.; Wang, M.; Arnason, J.; Mehta, A.; Purev, E.; Maloney, D.G.; Andreadis, C.; et al. Lisocabtagene maraleucel for patients with relapsed or refractory large B-cell lymphomas (TRANSCEND NHL 001): a multicentre seamless design study. Lancet 2020, 396, 839–852. 10.1016/S0140-6736(20)31366-0.

9. Sharma, P.; Kanapuru, B.; George, B.; Lin, X.; Xu, Z.; Bryan, W.W.; Pazdur, R.; Theoret, M.R. FDA Approval Summary: Idecabtagene Vicleucel for Relapsed or Refractory Multiple Myeloma. Clin. Cancer Res. 2022, 28, 1759–1764. 10.1158/1078-0432.CCR-21-3803.

10. Berdeja, J.G.; Madduri, D.; Usmani, S.Z.; Jakubowiak, A.; Agha, M.; Cohen, A.D.; Stewart, A.K.; Hari, P.; Htut, M.; Lesokhin, A.; et al. Ciltacabtagene autoleucel, a B-cell maturation antigen-directed chimeric antigen receptor T-cell therapy in patients with relapsed or refractory multiple myeloma (CARTITUDE-1): a phase 1b/2 open-label study. Lancet 2021, 398, 314–324. 10.1016/S0140-6736(21)00933-8.

11. Nagarsheth, N.; Wicha, M.S.; Zou, W. Chemokines in the cancer microenvironment and their relevance in cancer immunotherapy. Nat. Rev. Immunol. 2017, 17, 559–572. 10.1038/nri.2017.49.

12. Moon, E.K.; Wang, L.C.; Dolfi, D. V.; Wilson, C.B.; Ranganathan, R.; Sun, J.; Kapoor, V.; Scholler, J.; Puré, E.; Milone, M.C.; et al. Multifactorial T-cell hypofunction that is reversible can limit the efficacy of chimeric antigen receptor-transduced human T cells in solid tumors. Clin. Cancer Res. 2014, 20, 4262–4273. 10.1158/1078-0432.CCR-13-2627.

13. Nguyen, D.T.; Ogando-rivas, E.; Liu, R.; Wang, T.; Rubin, J.; Jin, L.; Tao, H.; Sawyer, W.W.; Mendez-gomez, H.R.; Cascio, M.; et al. CAR T cell Locomotion in Solid Tumor Microenvironment. Cells 2022, 1–26. 10.3390/cells11121974.

14. Ligon, J.A.; Wessel, K.M.; Shah, N.N.; Glod, J. Adoptive Cell Therapy in Pediatric and Young Adult Solid Tumors: Current Status and Future Directions. Front. Immunol. 2022, 13, 1–10. 10.3389/fimmu.2022.846346.

15. Majzner, R.G.; Ramakrishna, S.; Yeom, K.W.; Patel, S.; Chinnasamy, H.; Schultz, L.M.; Richards, R.M.; Jiang, L.; Barsan, V.; Mancusi, R.; et al. GD2-CAR T cell therapy for H3K27M-mutated diffuse midline gliomas. Nature 2022, 603, 934–941. 10.1038/s41586-022-04489-4.

16. Stadtmauer, E.A.; Fraietta, J.A.; Davis, M.M.; Cohen, A.D.; Weber, K.L.; Lancaster, E.; Mangan, P.A.; Kulikovskaya, I.; Gupta, M.; Chen, F.; et al. CRISPR-engineered T cells in patients with refractory cancer. Science (80-.). 2020, 367. 10.1126/science.aba7365.

17. Tobin, R.P.; Jordan, K.R.; Robinson, W.A.; Davis, D.; Borges, V.F.; Gonzalez, R.; Lewis, K.D.; McCarter, M.D. Targeting myeloid-derived suppressor cells using all-trans retinoic acid in melanoma patients treated with Ipilimumab. Int. Immunopharmacol. 2018, 63, 282–291. 10.1016/j.intimp.2018.08.007.

18. Zhang, W.; Moore, L.; Ji, P. Mouse models for cancer research. 2007, 149–152.

19. Kersten, K.; Visser, K.E.; Miltenburg, M.H.; Jonkers, J. Genetically engineered mouse models in oncology research and cancer medicine. EMBO Mol. Med. 2017, 9, 137–153. 10.15252/emmm.201606857.

20. Wu, Y.; Yu, X.Z. Modelling CAR-T therapy in humanized mice. EBioMedicine 2019, 40, 25–26. 10.1016/j.ebiom.2019.01.029.

21. Holzapfel, B.M.; Wagner, F.; Thibaudeau, L.; Levesque, J.P.; Hutmacher, D.W. Concise review: Humanized models of tumor immunology in the 21st century: Convergence of cancer research and tissue engineering. Stem Cells 2015, 33, 1696–1704. 10.1002/stem.1978.

22. Bareham, B.; Georgakopoulos, N.; Matas-Céspedes, A.; Curran, M.; Saeb-Parsy, K. Modeling human tumor-immune environments in vivo for the preclinical assessment of immunotherapies. Cancer Immunol. Immunother. 2021, 70, 2737–2750. 10.1007/s00262-021-02897-5.

23. Kleinman, H.K.; Martin, G.R. Matrigel: Basement membrane matrix with biological activity. Semin. Cancer Biol. 2005, 15, 378–386. 10.1016/j.semcancer.2005.05.004.

24. Aisenbrey, E.A.; Murphy, W.L. Synthetic alternatives to Matrigel. Nat. Rev. Mater. 2020, 5, 539–551. 10.1038/s41578-020-0199-8.

25. Caliari, S.R.; Burdick, J.A. A practical guide to hydrogels for cell culture. Nat. Methods 2016, 13, 405–414. 10.1038/nmeth.3839.

26. Tabdanov, E.D.; Rodríguez-Merced, N.J.; Cartagena-Rivera, A.X.; Puram, V. V.; Callaway, M.K.; Ensminger, E.A.; Pomeroy, E.J.; Yamamoto, K.; Lahr, W.S.; Webber, B.R.; et al. Engineering T cells to enhance 3D migration through structurally and mechanically complex tumor microenvironments. Nat. Commun. 2021, 12, 1–17. 10.1038/s41467-021-22985-5.

27. Ando, Y.; Siegler, E.L.; Ta, H.P.; Cinay, G.E.; Zhou, H.; Gorrell, K.A.; Au, H.; Jarvis, B.M.; Wang, P.; Shen, K. Evaluating CAR-T Cell Therapy in a Hypoxic 3D Tumor Model. Adv. Healthc. Mater. 2019, 8, 1–15. 10.1002/adhm.201900001.

28. Bhatia, S.N.; Ingber, D.E. Microfluidic organs-on-chips. Nat. Biotechnol. 2014, 32, 760–772. 10.1038/nbt.2989.

29. Grunewald, L.; Lam, T.; Andersch, L.; Klaus, A.; Schwiebert, S.; Winkler, A.; Gauert, A.; Heeren-Hagemann, A.I.; Astrahantseff, K.; Klironomos, F.; et al. A Reproducible Bioprinted 3D Tumor Model Serves as a Preselection Tool for CAR T Cell Therapy Optimization. Front. Immunol. 2021, 12, 1–14. 10.3389/fimmu.2021.689697.

30. Bhattacharjee, T.; Gil, C.J.; Marshall, S.L.; Urueña, J.M.; O’Bryan, C.S.; Carstens, M.; Keselowsky, B.; Palmer, G.D.; Ghivizzani, S.; Gibbs, C.P.; et al. Liquid-like Solids Support Cells in 3D. ACS Biomater. Sci. Eng. 2016, 2, 1787–1795. 10.1021/acsbiomaterials.6b00218.

31. Jensen, C.; Teng, Y. Is It Time to Start Transitioning From 2D to 3D Cell Culture? Front. Mol. Biosci. 2020, 7, 1–15. 10.3389/fmolb.2020.00033.

32. Verjans, E.T.; Doijen, J.; Luyten, W.; Landuyt, B.; Schoofs, L. Three-dimensional cell culture models for anticancer drug screening: Worth the effort? J. Cell. Physiol. 2018, 233, 2993–3003. 10.1002/jcp.26052.

33. Yamada, K.M.; Sixt, M. Mechanisms of 3D cell migration. Nat. Rev. Mol. Cell Biol. 2019, 20, 738–752. 10.1038/s41580-019-0172-9.

34. Merker, M.; Pfirrmann, V.; Oelsner, S.; Fulda, S.; Klingebiel, T.; Wels, W.S.; Bader, P.; Rettinger, E. Generation and characterization of ErbB2-CAR-engineered cytokine-induced killer cells for the treatment of high-risk soft tissue sarcoma in children. Oncotarget 2017, 8, 66137–66153. 10.18632/oncotarget.19821.

35. Chen, Z.; Han, S.; Sanny, A.; Chan, D.L.K.; van Noort, D.; Lim, W.; Tan, A.H.M.; Park, S. 3D hanging spheroid plate for high-throughput CAR T cell cytotoxicity assay. J. Nanobiotechnology 2022, 20, 1–14. 10.1186/s12951-021-01213-8.

36. Dillard, P.; Köksal, H.; Inderberg, E.M.; Wälchli, S. A spheroid killing assay by CAR T cells. J. Vis. Exp. 2018, 2018, 1–7. 10.3791/58785.

37. Timonen, T.; Saksela, E. A simplified isotope release assay for cell-mediated cytotoxicity against anchorage dependent target cells. J. lmmunological Methods 1977, 18, 123–132.

38. Bhat, R.; Rommelaere, J. NK-cell-dependent killing of colon carcinoma cells is mediated by natural cytotoxicity receptors (NCRs) and stimulated by parvovirus infection of target cells. BMC Cancer 2013, 13. 10.1186/1471-2407-13-367.

39. Wang, X.; Scarfò, I.; Schmidts, A.; Toner, M.; Maus, M. V.; Irimia, D. Dynamic Profiling of Antitumor Activity of CAR T Cells Using Micropatterned Tumor Arrays. Adv. Sci. 2019, 6. 10.1002/advs.201901829.

40. Fischer, K.; Mackensen, A. The flow cytometric PKH-26 assay for the determination of T-cell mediated cytotoxic activity. Methods 2003, 31, 135–142. 10.1016/S1046-2023(03)00123-3.

41. Matta, H.; Gopalakrishnan, R.; Choi, S.; Prakash, R.; Natarajan, V.; Prins, R.; Gong, S.; Chitnis, S.D.; Kahn, M.; Han, X.; et al. Development and characterization of a novel luciferase based cytotoxicity assay. Sci. Rep. 2018, 8, 1–14. 10.1038/s41598-017-18606-1.

42. Olivo Pimentel, V.; Yaromina, A.; Marcus, D.; Dubois, L.J.; Lambin, P. A novel co-culture assay to assess anti-tumor CD8+ T cell cytotoxicity via luminescence and multicolor flow cytometry. J. Immunol. Methods 2020, 487. 10.1016/j.jim.2020.112899.

43. Wallstabe, L.; Göttlich, C.; Nelke, L.C.; Kühnemundt, J.; Schwarz, T.; Nerreter, T.; Einsele, H.; Walles, H.; Dandekar, G.; Nietzer, S.L.; et al. ROR1-CAR T cells are effective against lung and breast cancer in advanced microphysiologic 3D tumor models. JCI Insight 2019, 4, 1–14. 10.1172/jci.insight.126345.

44. Schnalzger, T.E.; Groot, M.H.; Zhang, C.; Mosa, M.H.; Michels, B.E.; Röder, J.; Darvishi, T.; Wels, W.S.; Farin, H.F. 3D model for CAR -mediated cytotoxicity using patient-derived colorectal cancer organoids. EMBO J. 2019, 38, 1–15. 10.15252/embj.2018100928.

45. Zaman, M.H.; Trapani, L.M.; Siemeski, A.; MacKellar, D.; Gong, H.; Kamm, R.D.; Wells, A.; Lauffenburger, D.A.; Matsudaira, P. Migration of tumor cells in 3D matrices is governed by matrix stiffness along with cell-matrix adhesion and proteolysis. Proc. Natl. Acad. Sci. U. S. A. 2006, 103, 10889–10894. 10.1073/pnas.0604460103.

46. Kuczek, D.E.; Larsen, A.M.H.; Thorseth, M.L.; Carretta, M.; Kalvisa, A.; Siersbæk, M.S.; Simões, A.M.C.; Roslind, A.; Engelholm, L.H.; Noessner, E.; et al. Collagen density regulates the activity of tumor-infiltrating T cells. J. Immunother. Cancer 2019, 7, 1–15. 10.1186/s40425-019-0556-6.

47. Liu, J.; Tan, Y.; Zhang, H.; Zhang, Y.; Xu, P.; Chen, J.; Poh, Y.C.; Tang, K.; Wang, N.; Huang, B. Soft fibrin gels promote selection and growth of tumorigenic cells. Nat. Mater. 2012, 11, 734–741. 10.1038/nmat3361.

48. Sterner, R.C.; Sterner, R.M. CAR-T cell therapy: current limitations and potential strategies. Blood Cancer J. 2021, 11. 10.1038/s41408-021-00459-7.

49. Shaffer, D.R.; Savoldo, B.; Yi, Z.; Chow, K.K.H.; Kakarla, S.; Spencer, D.M.; Dotti, G.; Wu, M.F.; Liu, H.; Kenney, S.; et al. T cells redirected against CD70 for the immunotherapy of CD70-positive malignancies. Blood 2011, 117, 4304–4314. 10.1182/blood-2010-04-278218.

50. Pahl, J.H.W.; Santos, S.J.; Kuijjer, M.L.; Boerman, G.H.; Sand, L.G.L.; Szuhai, K.; Cleton-Jansen, A.; Egeler, R.M.; Bové, J.V.M.G.; Schilham, M.W.; et al. Expression of the immune regulation antigen CD70 in osteosarcoma. Cancer Cell Int. 2015, 15. 10.1186/s12935-015-0181-5.

51. Jenny Hendriks; Loes A. Gravestein; Kiki Tesselaar; René A. W. van Lier; Ton N. M. Schumacher; Jannie Borst CD27 is required for generation and long-term maintenance of T cell immunity. Nat. Immunol. 2000, 1, 433–440.

52. Jin, L.; Ge, H.; Long, Y.; Yang, C.; Chang, Y.E.; Mu, L.; Sayour, E.J.; De Leon, G.; Wang, Q.J.; Yang, J.C.; et al. CD70, a novel target of CAR T-cell therapy for gliomas. Neuro. Oncol. 2018, 20, 55–65. 10.1093/neuonc/nox116.

53. Ge, H.; Mu, L.; Jin, L.; Yang, C.; Chang, Y.E.; Long, Y.; DeLeon, G.; Deleyrolle, L.; Mitchell, D.A.; Kubilis, P.S.; et al. Tumor associated CD70 expression is involved in promoting tumor migration and macrophage infiltration in GBM. Int. J. Cancer 2017, 141, 1434–1444. 10.1002/ijc.30830.

54. Jin, L.; Tao, H.; Karachi, A.; Long, Y.; Hou, A.Y.; Na, M.; Dyson, K.A.; Grippin, A.J.; Deleyrolle, L.P.; Zhang, W.; et al. CXCR1-or CXCR2-modified CAR T cells co-opt IL-8 for maximal antitumor efficacy in solid tumors. 1–13. 10.1038/s41467-019-11869-4.

55. Nguyen, D.T.; Famiglietti, J.E.; Smolchek, R.A.; Dupee, Z.; Diodati, N.; Pedro, D.I.; Urueña, J.M.; Schaller, M.A.; Sawyer, W.G. 3D In Vitro Platform for Cell and Explant Culture in Liquidlike Solids. Cells 2022, 1–15.

56. McGhee, A.J.; McGhee, E.O.; Famiglietti, J.E.; Sawyer, W.G. In situ 3D spatiotemporal measurement of soluble biomarkers in spheroid culture. Vitr. Model. 2022, 309–321. 10.1007/s44164-022-00037-6.

57. Chandrashekar, D.S.; Karthikeyan, S.K.; Korla, P.K.; Patel, H.; Shovon, A.R.; Athar, M.; Netto, G.J.; Qin, Z.S.; Kumar, S.; Manne, U.; et al. UALCAN: An update to the integrated cancer data analysis platform. Neoplasia (United States) 2022, 25, 18–27. 10.1016/j.neo.2022.01.001.

58. Chandrashekar, D.S.; Bashel, B.; Balasubramanya, S.A.H.; Creighton, C.J.; Ponce-Rodriguez, I.; Chakravarthi, B.V.S.K.; Varambally, S. UALCAN: A Portal for Facilitating Tumor Subgroup Gene Expression and Survival Analyses. Neoplasia (United States) 2017, 19, 649–658. 10.1016/j.neo.2017.05.002.

59. Uhlen, M.; Zhang, C.; Lee, S.; Sjöstedt, E.; Fagerberg, L.; Bidkhori, G.; Benfeitas, R.; Arif, M.; Liu, Z.; Edfors, F.; et al. A pathology atlas of the human cancer transcriptome. Science (80-.). 2017, 357. 10.1126/science.aan2507.

60. Alam, O. A single-cell-type transcriptomics map of human tissues. Nat. Genet. 2021, 53, 1275. 10.1038/s41588-021-00938-4.

61. Sjöstedt, E.; Zhong, W.; Fagerberg, L.; Karlsson, M.; Mitsios, N.; Adori, C.; Oksvold, P.; Edfors, F.; Limiszewska, A.; Hikmet, F.; et al. An atlas of the protein-coding genes in the human, pig, and mouse brain. Science 2020, 367. 10.1126/science.aay5947.

62. Uhlén, M.; Fagerberg, L.; Hallström, B.M.; Lindskog, C.; Oksvold, P.; Mardinoglu, A.; Sivertsson, Å.; Kampf, C.; Sjöstedt, E.; Asplund, A.; et al. Tissue-based map of the human proteome. Science (80-.). 2015, 347. 10.1126/science.1260419.

63. Wang, Q.J.; Yu, Z.; Hanada, K.I.; Patel, K.; Kleiner, D.; Restifo, N.P.; Yang, J.C. Preclinical evaluation of chimeric antigen receptors targeting CD70-expressing cancers. Clin. Cancer Res. 2017, 23, 2267–2276. 10.1158/1078-0432.CCR-16-1421.

64. Stringer, C.; Wang, T.; Michaelos, M.; Pachitariu, M. Cellpose: a generalist algorithm for cellular segmentation. Nat. Methods 2021, 18, 100–106. 10.1038/s41592-020-01018-x.

65. Jacob, F.; Ming, G. li; Song, H. Generation and biobanking of patient-derived glioblastoma organoids and their application in CAR T cell testing. Nat. Protoc. 2020, 15, 4000–4033. 10.1038/s41596-020-0402-9.

66. Cazaux, M.; Grandjean, C.L.; Lemaître, F.; Garcia, Z.; Beck, R.J.; Milo, I.; Postat, J.; Beltman, J. B.; Cheadle, E.J.; Bousso, P. Single-cell imaging of CAR T cell activity in vivo reveals extensive functional and anatomical heterogeneity. J. Exp. Med. 2019, 216, 1038–1049. 10.1084/jem.20182375.

67. Uehara, S.; Grinberg, A.; Farber, J.M.; Love, P.E. A Role for CCR9 in T Lymphocyte Development and Migration. J. Immunol. 2002, 168, 2811–2819. 10.4049/jimmunol.168.6.2811.

68. Lança, T.; Costa, M.F.; Gonçalves-Sousa, N.; Rei, M.; Grosso, A.R.; Penido, C.; Silva-Santos, B. Protective Role of the Inflammatory CCR2/CCL2 Chemokine Pathway through Recruitment of Type 1 Cytotoxic γδ T Lymphocytes to Tumor Beds. J. Immunol. 2013, 190, 6673–6680. 10.4049/jimmunol.1300434.

69. Contento, R.L.; Molon, B.; Boularan, C.; Pozzan, T.; Manes, S.; Marullo, S.; Viola, A. CXCR4-CCR5: A couple modulating T cell functions. Proc. Natl. Acad. Sci. U. S. A. 2008, 105, 10101–10106. 10.1073/pnas.0804286105.

70. Barros, L.R.C.; Paixão, E.A.; Valli, A.M.P.; Naozuka, G.T.; Fassoni, A.C.; Almeida, R.C. CART math — A Mathematical Model of CAR-T Immunotherapy in Preclinical Studies of Hematological Cancers. 2021, 1–22.

71. Sahoo, P.; Yang, X.; Abler, D.; Maestrini, D.; Adhikarla, V.; Frankhouser, D.; Cho, H.; Machuca, V.; Wang, D.; Barish, M.; et al. Mathematical deconvolution of CAR T-cell proliferation and exhaustion from real-time killing assay data. J. R. Soc. Interface 2020, 17. 10.1098/rsif.2019.0734.

72. De Pillis, L.G.; Radunskaya, A.E.; Wiseman, C.L. A validated mathematical model of cell-mediated immune response to tumor growth. Cancer Res. 2005, 65, 7950–7958. 10.1158/0008-5472.CAN-05-0564.

73. Marofi, F.; Motavalli, R.; Safonov, V.A.; Thangavelu, L.; Yumashev, A.V.; Alexander, M.; Shomali, N.; Chartrand, M.S.; Pathak, Y.; Jarahian, M.; et al. CAR T cells in solid tumors: challenges and opportunities. Stem Cell Res. Ther. 2021, 12, 1–16. 10.1186/s13287-020-02128-1.

74. Caruana, I.; Savoldo, B.; Hoyos, V.; Weber, G.; Liu, H.; Kim, E.S.; Ittmann, M.M.; Marchetti, D.; Dotti, G. Heparanase promotes tumor infiltration and antitumor activity of CAR-redirected T lymphocytes. Nat. Med. 2015, 21, 524–529. 10.1038/nm.3833.

75. Friedl, P.; Entschladen, F.; Conrad, C.; Niggemann, B.; Zänker, K.S. CD4+ T lymphocytes migrating in three-dimensional collagen lattices lack focal adhesions and utilize ß1 integrin-independent strategies for polarization, interaction with collagen fibers and locomotion. Eur. J. Immunol. 1998, 28, 2331–2343. 10.1002/(SICI)1521-4141(199808)28:08<2331::AID-IMMU2331>3.0.CO;2-C.

76. Wolf, K.; Müller, R.; Borgmann, S.; Bröcker, E.B.; Friedl, P. Amoeboid shape change and contact guidance: T-lymphocyte crawling through fibrillar collagen is independent of matrix remodeling by MMPs and other proteases. Blood 2003, 102, 3262–3269. 10.1182/blood-2002-12-3791.

77. Paluch, E.K.; Aspalter, I.M.; Sixt, M. Focal Adhesion-Independent Cell Migration. Annu. Rev. Cell Dev. Biol. 2016, 32, 469–490. 10.1146/annurev-cellbio-111315-125341.

78. González-Martín, A.; Gómez, L.; Lustgarten, J.; Mira, E.; Mañes, S. Maximal T cell-mediated antitumor responses rely upon CCR5 expression in both CD4+ and CD8+ T cells. Cancer Res. 2011, 71, 5455–5466. 10.1158/0008-5472.CAN-11-1687.

79. Groom, J.R.; Luster, A.D. CXCR3 in T cell function. Exp. Cell Res. 2011, 317, 620–631. 10.1016/j.yexcr.2010.12.017.

80. Karin, N. CXCR3 Ligands in Cancer and Autoimmunity, Chemoattraction of Effector T Cells, and Beyond. Front. Immunol. 2020, 11, 1–9. 10.3389/fimmu.2020.00976.

81. Sautès-Fridman, C.; Petitprez, F.; Calderaro, J.; Fridman, W.H. Tertiary lymphoid structures in the era of cancer immunotherapy. Nat. Rev. Cancer 2019, 19, 307–325. 10.1038/s41568-019-0144-6.

82. Schumacher, T.N.; Thommen, D.S. Tertiary lymphoid structures in cancer. Science (80-.). 2022, 375. 10.1126/science.abf9419.

83. Munoz-Erazo, L.; Rhodes, J.L.; Marion, V.C.; Kemp, R.A. Tertiary lymphoid structures in cancer – considerations for patient prognosis. Cell. Mol. Immunol. 2020, 17, 570–575. 10.1038/s41423-020-0457-0.

84. Moreno-Smith, M.; Lutgendorf, S.K.; Sood, A.K. Impact of stress on cancer metastasis. Futur. Oncol. 2010, 6, 1863–1881. 10.2217/fon.10.142.

85. Adashek, J.J.; Subbiah, I.M.; Matos, I.; Garralda, E.; Menta, A.K.; Ganeshan, D.M.; Subbiah, V. Hyperprogression and Immunotherapy: Fact, Fiction, or Alternative Fact? Trends in Cancer 2020, 6, 181–191. 10.1016/j.trecan.2020.01.005.

86. Wildes, T.J.; Dyson, K.A.; Francis, C.; Wummer, B.; Yang, C.; Yegorov, O.; Shin, D.; Grippin, A.; DiVita Dean, B.; Abraham, R.; et al. Immune escape after adoptive T-cell therapy for malignant gliomas. Clin. Cancer Res. 2021, 26, 5689–5700. 10.1158/1078-0432.CCR-20-1065.

87. Stefanski, H.; Eaton, A.; Baggott, C.; Rossoff, J.; Verneris, M.R.; Keating, A.K.; Prabhu, S.; Pacenta, H.L.; Phillips, C.L.; Talano, J.-A.; et al. Higher doses of tisagenlecleucel associate with improved outcomes: a report from the pediatric real-world CAR consortium. Blood Adv. 2022, 6. 10.1182/bloodadvances.2022007246.

88. Keller, A.M.; Xiao, Y.; Peperzak, V.; Naik, S.H.; Borst, J. Costimulatory ligand CD70 allows induction of CD8+ T-cell immunity by immature dendritic cells in a vaccination setting. Blood 2009, 113, 5167–5175. 10.1182/blood-2008-03-148007.

